# Long read metagenomics, the next step?

**DOI:** 10.1101/2020.11.11.378109

**Authors:** Jose M. Haro-Moreno, Mario López-Pérez, Francisco Rodríguez-Valera

## Abstract

**Background:** Third-generation sequencing has penetrated little in metagenomics due to the high error rate and dependence for assembly on short-read designed bioinformatics. However, 2nd generation sequencing metagenomics (mostly Illumina) suffers from limitations, particularly in allowing assembly of microbes with high microdiversity or retrieving the flexible (adaptive) compartment of prokaryotic genomes.

**Results:** Here we have used different 3rd generation techniques to study the metagenome of a well-known marine sample from the mixed epipelagic water column of the winter Mediterranean. We have compared Oxford Nanopore and PacBio last generation technologies with the classical approach using Illumina short reads followed by assembly. PacBio Sequel II CCS appears particularly suitable for cellular metagenomics due to its low error rate. Long reads allow efficient direct retrieval of complete genes (473M/Tb) and operons before assembly, facilitating annotation and compensates the limitations of short reads or short-read assemblies. MetaSPAdes was the most appropriate assembly program when used in combination with short reads. The assemblies of the long reads allow also the reconstruction of much more complete metagenome-assembled genomes, even from microbes with high microdiversity. The flexible genome of reconstructed MAGs is much more complete and allows rescuing more adaptive genes.

**Conclusions:** For most applications of metagenomics, from community structure analysis to ecosystem functioning, long-reads should be applied whenever possible. Particularly for in-silico screening of biotechnologically useful genes, or population genomics, long-read metagenomics appears presently as a very fruitful approach and can be used from raw reads, before a computing-demanding (and potentially artefactual) assembly step.

## BACKGROUND

Metagenomics is still among the most powerful tools of exploratory Microbiology. Its application to several environments has allowed enlarging enormously what we know about the real (and largely unexpected) diversity of prokaryotic cells [1,2]. In actuality, these advances were largely possible by the advent of Illumina, high-throughput low-error short-read (SR) sequencing that has allowed enormous datasets that can be used for assembly of mock genomes called metagenome-assembled genomes (MAGs) [3,4]. Often supported by, typically incomplete, and expensive to generate, but largely reliable, single-cell amplified genomes (SAGs) [5]. MAGs have allowed rewriting most of what we knew about microbes during the last 10 years [6]. However, assembly driven metagenomics has weaknesses, i) low recovery of high micro-diversity microbes [7], ii) low recovery of the flexible genome [8] and iii) uncertainty due to potential chimera generation [9].

Long-read (LR) sequencing (i.e. Oxford Nanopore technologies –Nanopore, and Pacific Biosciences – PacBio) [10,11] solves major problems for genome assembly by covering large genomic tracks including the short to medium size repeats that mystify Illumina assembly algorithms [12–14]. Thus, it allows extremely fast and accurate closing of viral, prokaryotic, or even eukaryotic genomes [15–18]. However, these techniques are in general much more prone to error than Illumina what complicates their application for metagenomics. Therefore, high coverage of a homogeneous sequence is a must to get a reasonably reliable sequence [19]. However, the recent development of PacBio Sequel II chemistry allows decreasing enormously the error rate [20]. LR sequencing has the potential of fixing the problems of SR assembly and it also offers a good complementarity to SAGs since it is not biased by an amplification step and is simpler and much cheaper. LR metagenomes can be annotated directly from the sequence output avoiding erroneous protein translation and call. This would allow a good metabolic reconstruction of the environment with a high-accuracy prediction of biochemical activities. The core genome, the part best reconstructed in MAGs, is often the least interesting for ecological/biotechnological applications but could be reconstructed and exploited using LR. High reliable taxonomic affiliation by consensus similarity of multiple genes to a reference genome allows better inference of the origin of individual genes. Taxonomy markers such as ribosomal RNA operons can be retrieved complete allowing a reliable community structure determination [21]. To assess the resolving power of the two most advanced LR sequencing available (Oxford Nanopore and PacBio Sequel II) and compare it with Illumina we have selected a real metagenome rather than artificial genome mixtures [22–24] or low diversity environments [25,26]. We still do not know the real extent of the diversity of a real-life complex community to be able to mimic it with mixtures of known genomes. Besides, this kind of test have already been done and provided satisfactory results [22–24]. The open ocean is one of the oldest and most important communities for the global ecology of the planet and has been extensively studied by several methods, including metagenomics, for decades [27–30]. We took a sample from offshore Mediterranean waters in winter, when the water column is mixed, and it is likely that any depth sampled would provide a richer representation of the whole epipelagic layer microbiome [31]. From the same kind of sample, we have abundant information from previous metagenomic analysis [31–34]. We applied the two 3^rd^ generation sequencing platforms and analyzed the results pre and post assembly. We propose a specific pipeline based on CCS processing of the raw PacBio reads and their ulterior assembly to retrieve more useful information directly from the individual LRs, and much more complete MAGs than SR might allow.

## RESULTS AND DISCUSSION

### LR platforms output

A comparison of the metagenomic datasets generated by the three sequencing platforms is shown in **Table 1**. It is already apparent that with equivalent costs, one PacBio run produced 18 and 320 times more raw data (Gb) than Illumina or Nanopore sequencing, respectively. In addition, the largest sequenced read of ca. 448.5 Kb was also achieved with PacBio, whereas Nanopore produced the longest read of 35.5 Kb. However, when comparing the average read size, numbers were not so different, with 5.4 and 4.1 Kb for PacBio and Nanopore, respectively, i.e. extra long-reads are infrequent. Nanopore raw reads are known to have a significant error-rate in their sequences. Inspection of the dataset showed a variation comprised between 80 and 97 % read accuracy (**Figure S1**). Actually, 58% of the total sequence generated (851.5 Mb) had an error rate lower than 10% and only 98 Mb lower than 5 %. Also, we detected a shift in the GC content of the corrected sequences compared to the raw reads of all the other three methods (**Figure S2**) (28 %), indicating that error-correction in Nanopore can significantly alter the relative abundance of the prokaryotic taxa present in the sample. PacBio Sequel II does not provide the phred quality-score [35] of the dataset [36]. Therefore, comparisons at this level are not feasible.

**Table 1.** Summary statistics of the short-read and long-read sequencing technologies and protein-encoded genes retrieved from reads.

| Sequencing Technology | Illumina<br>(Nextseq<br>2x100bp) | Oxford Nanopore (Minlon<br>Flow Cell, v9.4R) |  | PacBio Sequel II (8M SMRT Cell Run) |  |  |  |
| --- | --- | --- | --- | --- | --- | --- | --- |
| Read Type (Proccesing) | Trimmed<br>Reads | Raw<br>Reads | Corrected Reads | Raw Reads | CCS5 | CCS10 | CCS15 |
| Sequencing statistics: |  |  |  |  |  |  |  |
| #Sequences (Millions) | 234.5 | 0.3 | 0.1 | 81.4 | 2.8 | 1.9 | 1.5 |
| #Nucleotides sequenced (Gb) | 23.4 | 1.4 | 0.4 | 439.6 | 15.4 | 10.1 | 7.6 |
| Largest read size (bp) | 100 | 35,490 | 22,220 | 448,515 | 36,542 | 24,735 | 17,976 |
| Average length read size (bp) | 99.6 | 4,117.9 | 3,148.2 | 5,401.6 | 5,422.4 | 5,190.7 | 4,968.8 |
| Predicted proteins (Reads > 1 Kb): |  |  |  |  |  |  |  |
| #Proteins (Millions) | -- | 1.3 | 1.5 | 368.1 | 21.6 | 12.2 | 9.0 |
| Average Protein Size (aa) | -- | 85.9 | 83.2 | 90.4 | 195.1 | 241.5 | 248.4 |
| Proteins / Mb Sequenced | -- | 937.3 | 3,907.9 | 837.3 | 1,407.1 | 1,212.8 | 1,184.4 |

However, to guaranty a low error rate of PacBio, we applied the software “Highly Accurate Single-Molecule Consensus Reads” (CCS reads) [17]. The algorithm selects DNA tracts that are re-sequenced up to the number provided (5, 10, or 15 times). These numbers theoretically achieve 99, 99.9 and 99.95 % base call accuracy, respectively. Thus, for example, the total PacBio sequence generated decreased from 439.63 Gb (raw) to 7.63 Gb (CCS15) (**Table 1**). To assess the read accuracy, we assumed that erroneous nucleotides would lead to an increase of stop codons in the predicted proteins and could be measured by the decrease of their average protein size. As seen in **Table 1** PacBio raw reads had error rates similar to those of Nanopore (average protein size 90.4 and 85.9 amino acids respectively) while CCS15 provided an average protein size of 248.4 amino acids, much closer to the values of the two dominant microbes in these waters (302.5 or 255 found for *Ca*. Pelagibacter HTCC7211 and *Prochlorococcus marinus* MED4 pure culture genomes, respectively). Thus, we have concluded that the quality of CCS15 is enough to get a reliable picture of the genes present in the sample. Contrastingly, Nanopore does not seem suitable for cellular (as opposed to viral [18,37]) metagenomic analysis. Henceforth, we will focus the analysis on the PacBio results that we will refer to as long-reads (LR) and Illumina Nextseq as short-reads (SR).

### Taxonomic profiling of samples by metagenomic rRNA operons

The community structure of a metagenomic sample is one of the most basic pieces of information about a microbial assemblage and can be assessed by multiple approaches [38,39]. One of the most reliable is the isolation of SRs that have hits to 16S rRNA genes and use their huge databases to affiliate the sequences and thus the microbes behind. This can be done with the individual SRs or with SR assembled rRNA genes [40], although assembly of these highly conserved sequences is not very reliable. In the case of LR sequencing, complete (or nearly so) rRNA genes and even operons can be retrieved within a single contig making assembly superfluous [21,41]. We have extracted and compared 16S rRNA gene fragments to check whether LRs can improve the taxonomic affiliation. Thus, we were able to extract 9,763 16S rRNA sequences from LR CCS15 of our sample (average length: 1,207 bp; 0.34 % of total LRs), and 20,564 SRs (average length: 95 bp; 0.22% of total SRs). These sequences were classified against the SILVA database (**Table S1**), The community structure derived from LRs was nearly identical to the one obtained from SRs down the level of families (**Figure 1A and Table S1**), with only a significant exception in Cyanobacteria, that were overrepresented in the SR dataset, 9.1 % compared to 6.9 % in the LRs. However, the availably of longer gene fragments with LRs improves the 16S rRNA classification, decreasing the number of reads that were not classified to any specific phylum (0.4 % LR versus 1.3 % SR,) or could not be ascribed to lower-level taxa (4.3 % that only reached the class level Alphaproteobacteria with SRs versus only 1.3 % LRs), (**Figure 1A and Table S1**). These results indicate a better resolution microbial classification of LRs. More importantly, LRs have the potential to uncover complete 16S rRNA sequences from “dark matter” [2] microbes with a higher level of resolution and reliability avoiding potential assembly artifacts.

**Figure 1.**
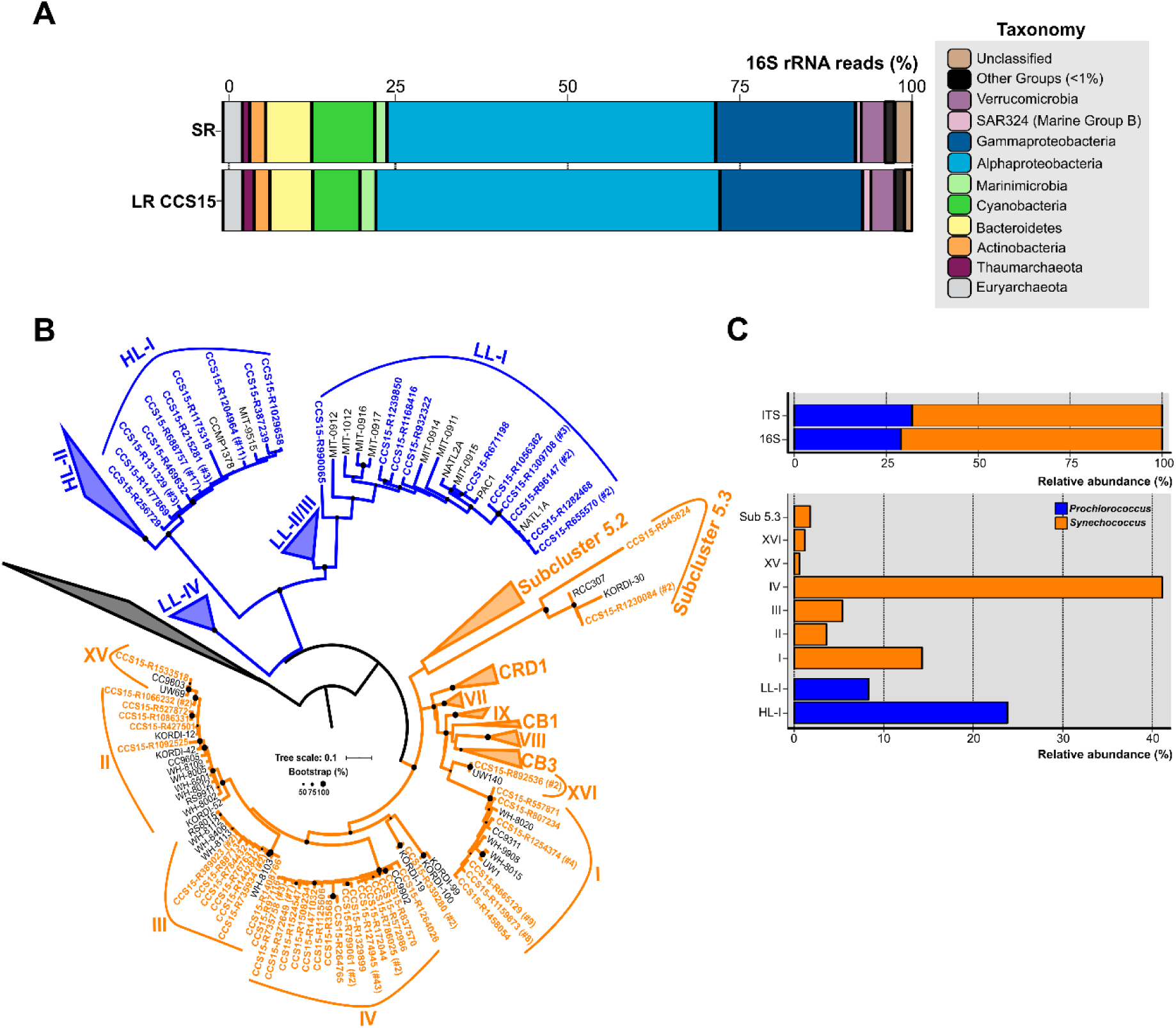
**A**. Phylum-level composition based on 16S rRNA gene fragments of Illumina (SR) and PacBio CCS15 metagenomic reads. The phylum Proteobacteria was divided into its class-level classification. Only those groups with abundance values larger than 1% are shown. **B**. Maximum-likelihood phylogenetic tree of the 16S-23S rRNA genes ITS extracted from CCS15 reads classified by 16S rRNA as Cyanobacteria. Sequences are coloured according to their affiliation to *Prochlorococcus* (blue) or *Synechococcus* (orange). In order to simplify the tree, nucleotide sequences were dereplicated at 99% identity. Numbers between brackets indicate the number of sequences that clustered at this level to a given CCS15 read (in bold). ITS from reference isolates are also shown (black). **C**. Upper panel, relative abundance of *Synechococcus* and *Prochlorococcus* identified from ITS or 16S rRNA sequences in the CCS15 reads. Bottom panel, ITS classified into subclades and ecotypes.

Furthermore, other useful identifiers within the ribosomal operon, including hypervariable regions such as the internal transcribed spacers (ITS), could be retrieved within a single contig [42,43]. These allow a precise community structure determination that includes ecotypes or even strains. As an example, the ITS tree for the picocyanobacterial retrieved in our sample is shown in **Figure 1B**. We considered only complete ITS sequences (both 16S and 23S genes had to be present in the same read). 170 ITSs could be extracted, of which 68 % were classified as *Synechococcus*. Within this genus, clades IV (69 ITS) and I (24 ITS) were the most dominant in the sample (**Figure 1C**). These clades have been detected before in cold coastal waters [44,45] so their presence in our mixed winter sample was expected. Along similar lines, two *Prochlorococcus* ecotypes dominated the sample. Forty out of 54 ITSs were assigned to the High-Light I (HL-I) ecotype, while only 14 sequences grouped within the Low-Light I (LL-I) ecotype (**Figure 1C**). These results fit well with genome recruitment data using pure cultures or MAGs of the different ecotypes as reference carried out on in this and similar samples collected different years, seasons, and depths [31,32], supporting the reliability of the LR ITS data.

### Metagenomic assembly with LR

Still, the possibility to retrieve complete (or nearly so) genomes from metagenomes (MAGs) is highly informative for understanding uncultivated microbes. In principle, the application of LR to a complex sample could improve metagenomic assembly by simplifying the leap across repeats that hamper SRs assembly. However, the choice of an assembler for metagenomic projects is not trivial. To gauge the applicability of different programs we have to consider also the possibility of a hybrid assembly to take advantage of the high coverage and low error rate of SRs. We selected two specific assemblers for LRs (based on overlaps, Canu or Bruijn Graphs, MetaFlye) and one that is hybrid and can combine SRs and LRs (see methods). Only assembled contigs larger than 5 Kb have been further considered.

In a first approach, five subsets of LRs (before CCS processing) larger than 7Kb were assembled (**Table S2**). Assembly results by metaFlye and Canu were positively correlated (close to linear, data not shown) with the sequencing effort. The largest contig size also followed this pattern, while the average contig size was not variable within the range considered. Besides, these assemblers resulted in a low number of proteins per Mb and small average protein size, indicating low-quality assembly (**Table S2**). Conversely, metaSPAdes (hybrid assembly LR and SR) showed that the effect of assembling larger amounts of PacBio raw reads did not have a linear trend in the resulting assembly. Besides, due to the inclusion of SRs in the assembly process, at lower coverage values, metaSPAdes assembly was larger than the other two assemblers. Furthermore, given the restricted use of LRs in the hybrid assembly (it mainly comes from the SRs), the high error rate of LRs did not affect the quality (average size) of the resulting assembled proteins (**Table S2**). Thus, metaSPAdes appears as the best option for assembly of LR as long as an SR dataset is also available.

We also evaluated the effect of increasing steps of CCS in the assembly by the three software packages compared to SRs IDBA assembly (SRa). Regardless of the CCS steps (5, 10, or 15), the resulting assembly outperformed IDBA SR (**Figure 2A**). The largest contig was achieved with metaSPAdes CCS5, 2.6 Mb long, one order of magnitude higher than the one achieved with SRa (275 Kb). Besides, the average genome size was also seven times higher with metaSPAdes CCS5 (**Figure 2A**). Although it yielded a lower assembled output than the other two methods, the contigs had smaller average protein sizes (**Figure 2B lower panel**) nullifying the advantage. Therefore, the best assembly results in terms of assembly size and, particularly, reliability were achieved using metaSPAdes with the pool of CCS15 LRs (**Figure 2A and B**). Unfortunately, we lose the longest fragments obtained in the CCS5 assembly, likely due to a decrease in the coverage (**Table 1**). To validate the assembly of metaSPAdes CCS15 (**Figure 2C**) we have compared the large-scale taxonomic affiliation of the contigs with those of the SRa (**Figure 2C**). All phyla were recovered (and in similar proportions) by both methods and only small numerical differences were found confirming that no major bias (at least not different from those that might possess SRa) was acting on the retrieval of microbes by metaSPAdes CCS15 (henceforth LRa).

**Figure 2.**
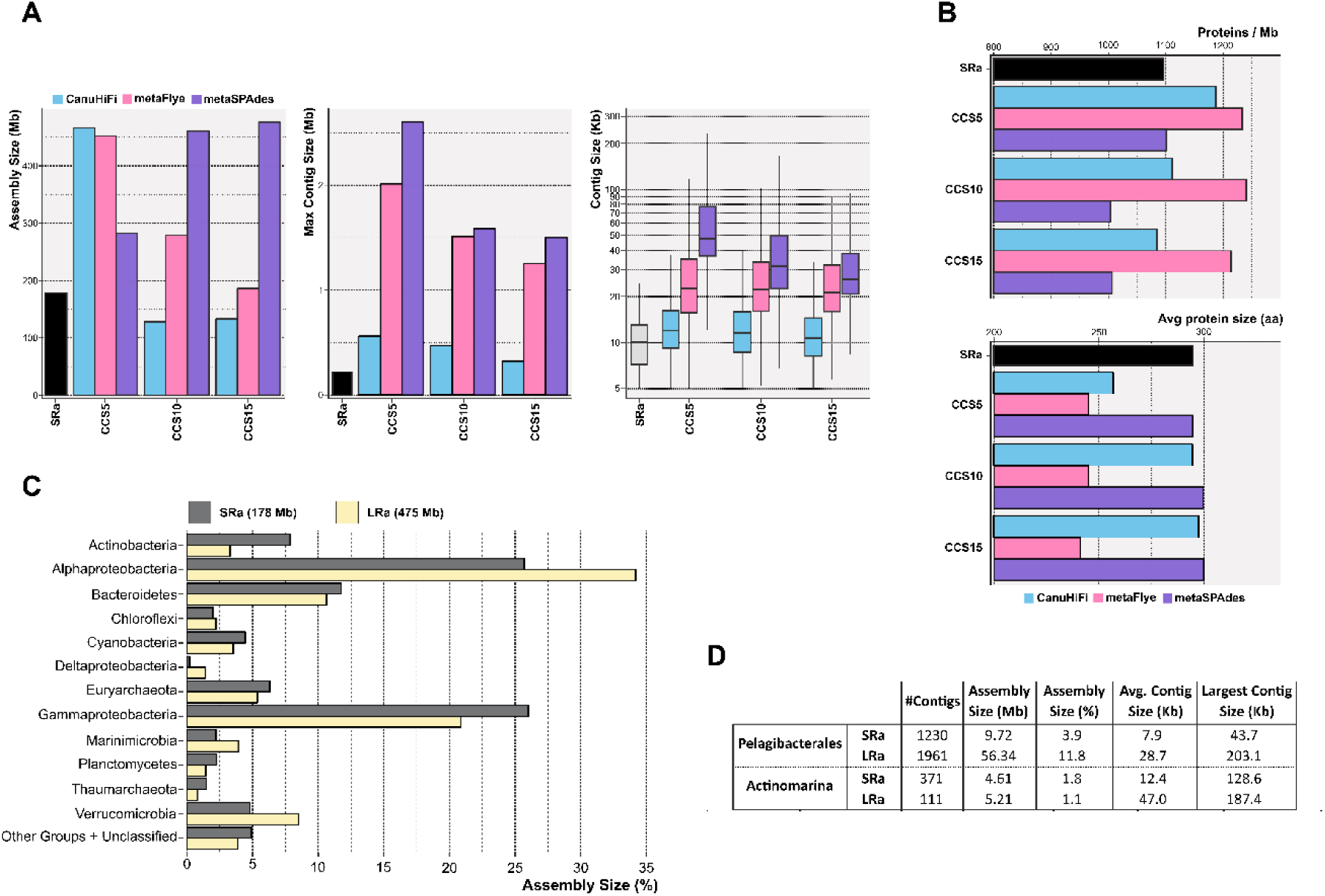
**A**. Bar and box plots indicating the total assembly size, maximum contig length and contig size distribution for the Illumina (SR) and PacBio CCS5, CCS10 and CCS15 assemblies. CCS reads were assembled with CanuHiFi, metaFlye and metaSPAdes (blue, pink and violet bars, respectively). **B**. Similar to A, but representing the number of predicted proteins assembled per Megabase (upper panel) and the average protein size (lower panel). **C**. Taxonomic classification at the level of phylum of the resulting Illumina (SRa) and PacBio CCS15 (LRa) assemblies. The phylum Proteobacteria was divided into its class-level classification. **D**. Summary of the assembly statistics for contigs classified as *Ca*. Pelagibacterales and *Ca*. Actinomarinales.

One major problem of the classical MAG approach is its proven low yield of some of the most prevalent members of the community. A very prominent example in the marine environment is the Pelagibacterales [46]. Despite their dominance in open epipelagic marine waters [46], the numbers of MAGs retrieved in metagenomic studies are relatively small, with only 34 MAGs (medium quality, >50% complete and < 5% contaminated) available presently in public repositories [7]. Another example is *Ca*. Actinomarinales [47], a cosmopolitan marine actinobacterium that for accounts up to 5% of the prokaryotic community and has only seven MAGs available [48]. The reasons for this anomaly are unclear but the most likely explanation points to the high level of sequence microdiversity characteristic of these microbes. Here, the use of LR metagenomics improved considerably the assembly of both microbes (**Figure 2D**). LRa achieved a better assembly size, as in Pelagibacterales, with ∼6 times more data with LRa than SRa, and longer contigs that might help the recovery of complete (or nearly so) MAGs.

### Recovery of novel genes

One of the major levers of metagenomics is its capacity to identify new proteins that alone can provide important insights into the ecology of the sample and often is capable to associate the activities to certain microbial groups. In any case, the expression in surrogate vectors, allows the use of the recovered proteins for structural or biotechnological studies. LRs can span complete genes (or operons) and, therefore, avoid the SR assembly step, which is greatly biased by the choice of the assembler and gene calling [49,50] together with the (micro)diversity and abundance of prokaryotes in the sample [51]. A recent study [52] evaluated how the application of high-throughput metagenomic sequencing has severely improved the catalog of marine microbial genes. Large metagenomic studies, such as *Tara* Oceans [30], GEOTRACES [53] and Malaspina [54], sequenced and assembled hundreds of marine SR datasets at different years, seasons, latitudes, and depths. They have retrieved ca. 50M non-redundant proteins [52]. Yet, when this number is normalized by the sequencing effort (4.8, 4.8 and 52.1M non-redundant proteins/Tb, respectively [52]), the numbers become smaller than those retrieved by the Global Ocean Sampling by cloning and Sanger sequencing (GOS [55]) (624 M /Tb [52]). In our work, LR sequencing of just one metagenomic sample yielded 3.6 M non-redundant proteins, what can be extrapolated to 473.7M/Tb, very close to the GOS numbers, but with a largely diminished cost/person-power investment, and better yield of reconstructed genomes (see below) and gene clusters. To assess further the differential capability to recover novel proteins by LR metagenomics we have selected to search in our single metagenomic sample for four common objectives of screenings for biotechnologically relevant proteins or gene clusters: rhodopsins, polyketide synthases (PKS), and CRISPR systems.

One of the best examples of the biotechnological harvest of metagenomics has been the retrieval of a vast diversity of retinal proteins (rhodopsins) [56–58] critical for the development of optogenetics, a technology with remarkable potential in neurobiology and medicine [59,60]. The photic zone of the ocean is the quintessential habitat to screen for the diversity of rhodopsins and already many have been retrieved by SR assembly metagenomics [61,62]. Remarkably, the largest numbers of rhodopsins (>200 amino acids, clustered at 90 % amino acid identity) were found in the individual LRa (330 rhodopsin genes/Gb assembled) (**Figure 3A)**. However, considering the sequencing effort to assemble (31 Gb, sum of SR and LR CCS15), this number decreases down to 5 rhodopsins / Gb, smaller than the LR output (50 rhodopsins / Gb). This result illustrates the performance of LR to recover novel proteins without the biased assembly step. Besides, a clustering at >30% identity of all the rhodopsins retrieved with LR CCS15 (#2858) and *Tara* assemblies (#5887) resulted in 25 distinct protein clusters (data not shown), where twelve of them grouped sequences from both datasets. Eleven clusters had only sequences originating from *Tara* assemblies. *Tara* samples span different locations, depths, and seasons; thus, it was to be expected that its dataset contained a higher diversity of rhodopsins than our single sample. Nonetheless, we could identify three novel rhodopsin clusters without *Tara* representatives, indicating there is still room for the discovery of novel rhodopsins through LR metagenomics. In fact, one of these three clusters contained sequences similar (67% amino acid identity) to RubyACRs [63], a recently reported anion channelrhodopsin with promising implications in optogenetics.

**Figure 3.**
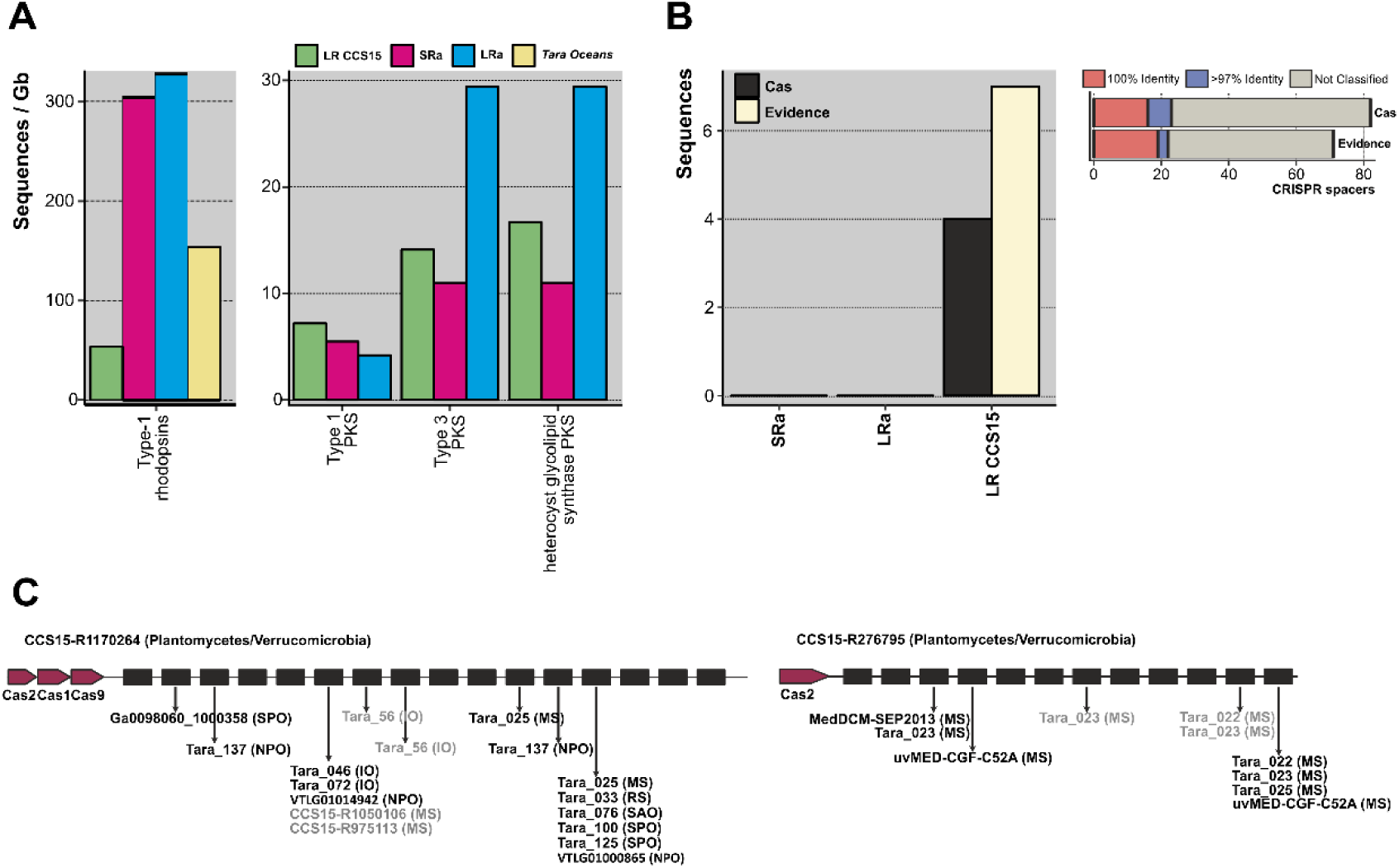
**A**. Number of type-1 rhodopsins and polyketide synthases (PKS) (left and right panels, respectively), retrieved from the LR CCS15 reads (green bar), from SRa and LRa assemblies (red and blue blars, respectively) and from *Tara Oceans* assemblies (yellow bar). Hits are normalized by the size of the database (in Gigabases). **B**. Left panel, number of sequences (LR CCS15, and SRa and LRa contigs) containing CRISPR and Cas proteins (coloured in black) or only a CRISPR array with an evidence score ≥ 4 (light yellow bar). Right panel, number of CRISPR spacers classified at 100 and >97% identity. Sequences failing the 97 % identity threshold or not matching to the database are grouped into the “not-classified” bar. **C**. Two examples of metagenomic reads having a CRISPR array and Cas proteins. Sequences in black and grey represent 100 and >97% identity hits. Between brackets the isolation source of the hit: Mediterranean Sea (MS); Red Sea (RS); Indian Sea (IO), North and South Atlantic Oceans (NAO, SAO); and North and South Pacific Oceans (NPO, SPO).

From a biotechnological point of view, one of the most important natural products is polyketide synthases [64,65]. They are large proteins and often require other accompanying genes to be functional. Besides, they tend to be located at the flexible genome that as mentioned before assembles poorly in SR metagenomes. The total number of PKS type 1 (long and modular proteins) was similar to the three datasets (LRs, SRa, and LRa). The other two types, type 3 (smaller proteins that work only with the complement of the other members of the cluster) and heterocyst glycolipid synthase PKS (cyanobacterial PKSs) were better recovered by LRa (**Figure 3A**). Actually, in LR individual reads there were type 1 complete clusters. One of them was 100% similar to the 1-heptadecene biosynthetic gene cluster from *Cyanothece* sp. PCC 7822 [66]. Some type 3 PKS (mostly chalcone synthases from *Synechococcus*), were also recovered complete (data not shown). Thus, as in the case of rhodopsins assembly seems rather redundant for LR screening for PKSs.

Although CRISPR systems are very scarce in seawater, these systems are also often screened for and described from metagenomic datasets [67]. They form large clusters of Cas (CRISPR associated) proteins together with long stretches of tandem repeats [68]. We could find four LRs containing both Cas proteins and complete CRISPR arrays (**Figure 3B**); this number increased up to 15 if we included sequences with no Cas proteins but with an evidence value ≥ 4 following CRISPRdetect [69]. A comprehensive search of the spacers in a custom database containing metagenomes, viromes, and reference viral sequences (see methods) showed that 28 % of them were positively affiliated to viral sequences (**Figure 3B**). LRs CCS15-R1170264 and CCS15-R276795 represent two CRISPR arrays that affiliated with two different and uncultured Planctomycetes/Verrucomicrobia bacteria (**Figure 3C**). However, their spacers indicated two different geographic distributions. Spacers in CCS15-R1170264 matched several sequences recovered from different locations, indicating a widespread distribution of the microbe. Conversely, CCS15-R276795 showed Mediterranean endemism, given that those spacers matched exclusively viral sequences recovered from metagenomes [30] and fosmid libraries [70,71] from that sea.

### Recovery of genomes

To assess the efficiency of MAG retrieval by LRa we extracted 77 MAGs (>50% completeness, >5% contamination). This figure is rather small compared to other similar studies carried out by SRa. However, when corrected for the amount of processed sequence (CCS15 only) the ratio is higher than in similar studies carried out with similar samples by SRa, as well as the average contig size and the degree of completeness (**Figure 4**). To compare the MAG reconstruction carried out by both approaches, we selected 31 MAGs retrieved in the previous SRa works carried out in our laboratory [31,34] and that had >99.5% average nucleotide identity (ANI) to MAGs recovered by LRa in this work (**Table S3**). LRa MAGs were on average 1.5 larger than the SRa MAGs, but even more remarkable, the largest contig by LRa was 2.7 larger and the average contig size was 4 times larger, what clearly turns the balance in favor of LRa to get high-quality reliable MAGs.

**Figure 4.**
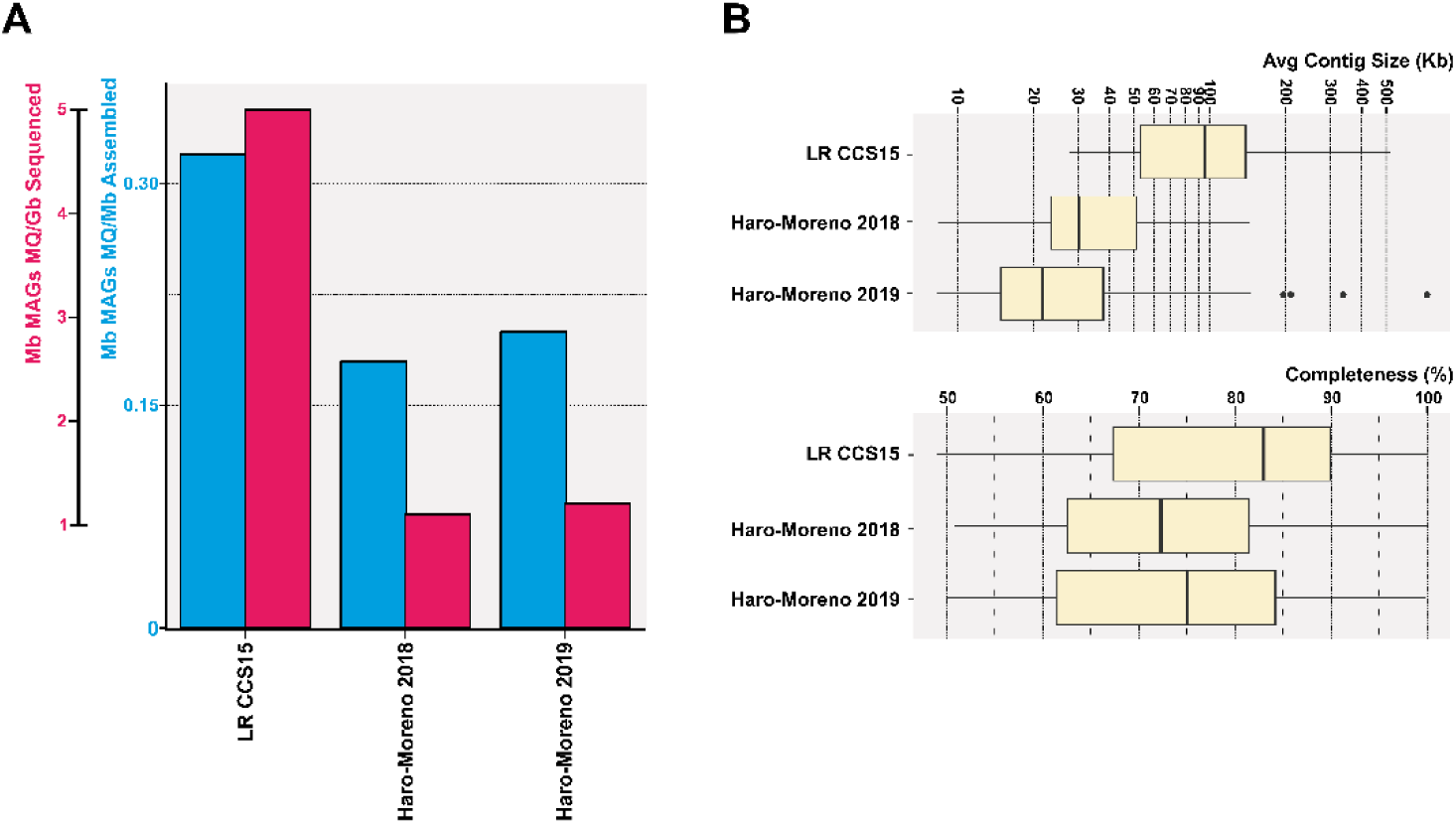
A. Contribution of medium-quality MAGs (>50% complete and <5% contamination) recovered in this study (LR CCS15) and from two previous reports from the same location in the Mediterranean Sea, Haro-Moreno *et al*. 2018 [31] and Haro-Moreno *et al*. 2019 [34]. Values are normalized by the assembly size and sequencing effort (blue and red bars, respectively). **B**. Box plots showing the average contig size and the degree of completeness of MAGs described in A.

Visual inspection of the LRa MAGs indicated very complete and easily closable collections of contigs. To objectively compare the completion of MAGs generated by both approaches we could identify two microbial genomes that are derived from pure cultures and were retrieved also in LRa and SRa, and compared the assemblies with the reference genomes (assumed to be complete) **(Figure 5)**. One of them was a genome similar (ANI 97%) to the cyanobacterium *Prochlorococcus marinus* MED4 (high-light-adapted ecotype) [72], one of the most abundant microbes in our kind of sample (30 RPKG in the sample analyzed here). The SRa was only 2% complete (estimated by CheckM) with 6 small contigs among which the longest was 34 kb (**Figure 5A**). The LRa MAG covered nearly the complete pure culture genome, with only eight contigs, the longest 608kb, and with more than 98% of the pure culture genome (**Figure 5A)**. Gaps were found mostly at the location of the known major flexible genomic island of this microbe [73] particularly, GI4 that codes for the O-chain polysaccharide [74]. We also reconstructed by both assemblies a relative (93.4% ANI) of the Thaumarchaeon *Ca*. Nitrosomarinus catalina SPOT01 [75] by LRa and SRa. LRa produced only three contigs, the longest being 1Mb with a 99% completeness based on checkM (data not shown). The comparison with the reference genome *Ca*. Nitrosomarinus catalina SPOT01 showed that two regions were not covered (**Figure 5A**). One largely corresponds to a prophage that might not be present in our local relative, and the other again a genomic island putatively involved in the synthesis of a polysaccharide equivalent to GI4 in MED4 [73].

**Figure 5.**
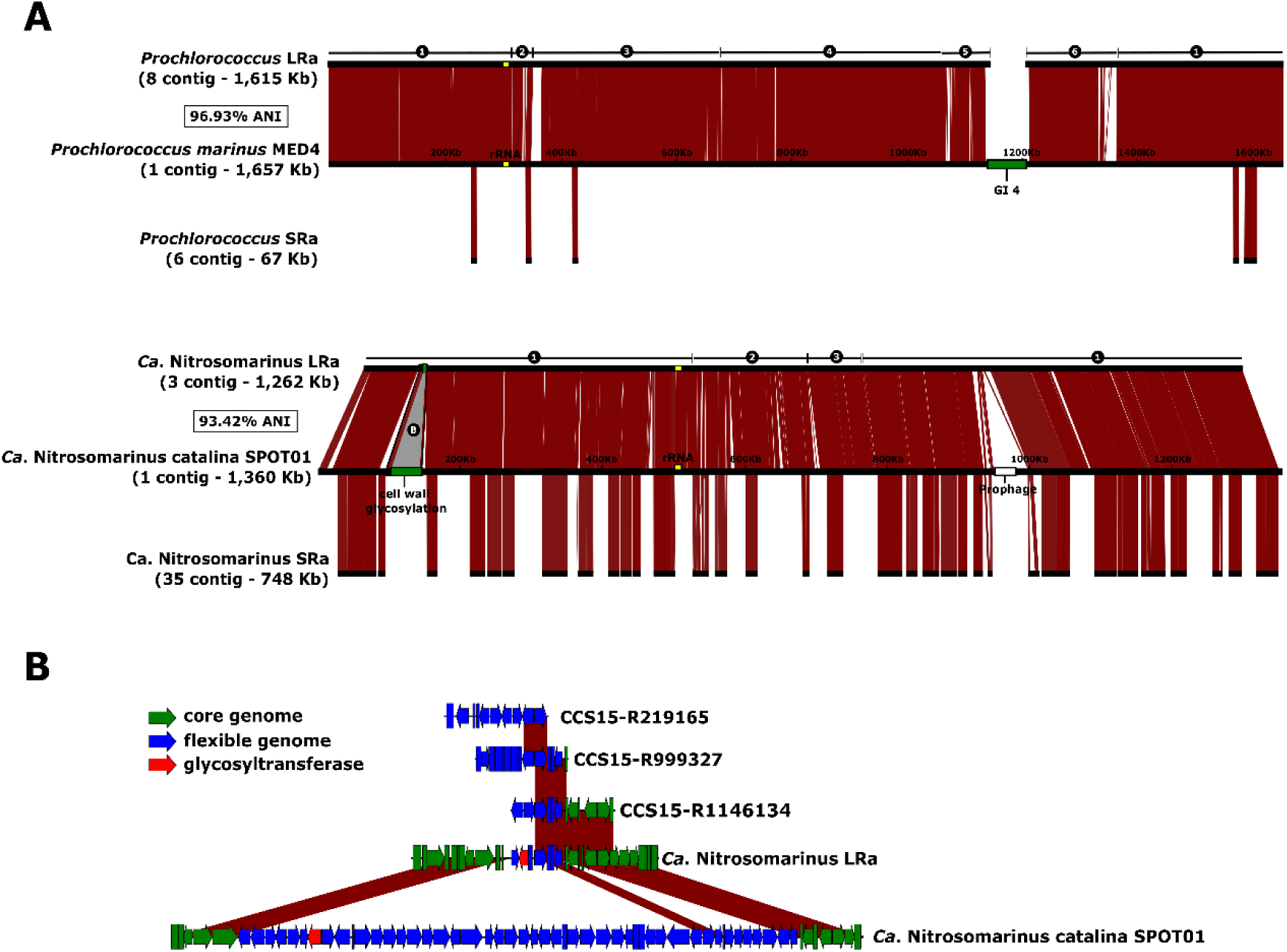
**A**. Alignment of the reference genomes *Prochlorococcus marinus* MED4 and *Ca*. Nitrosomarinus catalina SPOT01 against the reconstructed MAGs from LR CCS15 (above) and SR (below). Number of contigs, and ANI of the LR CCS15 MAGs and the reference genomes indicated to the left. GI4 of *P. marinus* coding for O-chain polysaccharide synthesis [74] and a putative cell was glycosylation cluster in of *Ca*. N. catalina are in green. **B**. Magnified B region as labelled in A indicating also some syntenic fragments found among CCS15 reads.

It has been established that there are two main categories of flexible genomic islands (fGIs) in prokaryotic genomes: i) replacement fGIs are involved in synthesizing the outer glycosidic envelope of the cells (such as the O-chain in Gram-negatives) [76] that varies between closely related strains, and ii) additive fGIs (such as integrons) that vary more gradually by replacement of relatively small gene cassettes that appear among other gene clusters conserved between strains. One of the problems inherent to the assembly of short reads is the failure to assemble fGIs in prokaryotic genomes. The reasons are multifactorial, i) SRa contigs belonging to fGIs tend to bin separately due to variations in genomic parameters, ii) they are less abundant since are only harbored by some lineages within the population, and finally iii) replacement fGIs tend to be surrounded by highly variable (if conserved) genes that are followed by totally divergent sequences [77]. All these scenarios make assembly algorithms highly inefficient in retrieving fGIs. This is a major setback since many genes of biotechnological potential are found within flexible genomic islands. Furthermore, many key ecological functions such as transporters, degradation of resilient compounds, virulence factors and many others are also found in these genomic regions [78,79]. That the long replacement GI4 do not assemble in the MED4 LRa MAG was to be expected given the high diversity of very polyclonal microbes such as *Prochlorococcus* [80] and the length of this specific fGI involved in the synthesis of the O-chain polysaccharide [81,82]. The presence of multiple (and long) versions of GI4 might disorient the assembler that has many possibilities to continue the contig. On the other hand, the small island present in the reconstructed *Ca*. Nitrosomarinus MAG (**Figure 5B)** might be short enough to appear in one single read and it did appear in the MAG and other identical fragments recovered as LRs.

To assess the improvement in the retrieval of flexible genomic islands by LRa, we in more detail one marker that is usually found in these islands: arrays of glycosyl transferases (GT). Genome analyses have demonstrated that in this GI there is an accumulation of GTs and, therefore, these genes are a good indicator for the recovery of such gene clusters. We have considered only fragments between 5 Kb encoding for at least 5 GTs in a window of maximum 20 genes as putative parts of these glycosylation islands. Indeed, LRa recovered more than 300 GT/Gb while SRa only found 100. The presence of more GTs in the LRa can be explained by the short stretches at the beginning of the island recovered at the end of the assembly. Additive flexible GIs that contain only small differential cassettes, with conserved clusters alternating with variable ones, that can be straddled by individual reads, would be recovered much more efficiently, which would explain the increase of typical additive GIs components such as the PKS or CRISPR (see above).

## CONCLUSIONS

### Is LR metagenomics the next step?

The answer appears to be largely affirmative. Although SR approaches might be used to complement for recruitment of known genomes or to improve assembly, for most purposes LR sequencing is much more rewarding both in terms of the amount and quality of the information. It only requires slightly more environmental DNA and of better quality (more care should be taken to avoid too much fragmentation of the DNA in the sample) and the cost per properly annotated gene is significantly lower. Furthermore, it allows a first glimpse at the flexible genome of many microbes in which a wealth of potentially useful biotechnology might be hidden. It might complement SAGs and MAGs to get complete and reliable genomes of the many novel groups that have been uncovered during the last decade, improve their annotation and their representation in databases and eventually lead to a more realistic picture of the real diversity of microbes. The enhanced recovery of the flexible genome would provide a better understanding of their ecological features and their potential applications. Last but not least, the intricacy of natural populations of bacteria could be analyzed in detail providing a glimpse at microbial evolution in action.

## MATERIAL AND METHODS

### Sampling, processing, and sequencing

Samples from two different depths (20 and 40 m) were collected on February 15^th^ 2019 from the epipelagic Mediterranean Sea, at 20 nautical miles off the coast of Alicante (Spain) (37.35361°N, 0.286194°W), during winter where the water column is mixed. This location has been studied previously by metagenomic approaches [31–34,70,83]. For each depth, 200 L were collected and filtered on board as described in [31]. Briefly, seawater samples were sequentially filtered through 20, 5 and 0.22 µm pore filter polycarbonate filters (Millipore). Water was directly pumped onto the series of filters to minimize the bottle effect. Filters were immediately frozen on dry ice and stored at -80 °C until processing.

DNA extraction was performed from the 0.22 µm filter (free-living bacteria) following the phenol:chloroform extraction. Given the large amount of DNA needed for sequencing, DNA from the two samples (20 and 40 m) was pooled together. Metagenomes were sequenced using Illumina Nextseq (100 bp, paired-end reads) (Macrogen, South Korea), and using Oxford Nanopore (one MinIon flowcell run, chemistry version 9.4R) and PacBio Sequel II (one 8M SMRT Cell Run, 30-hour movie) (Genomics Resource Center, University of Maryland, USA).

### Raw read filtering and assembly of metagenomic samples

The quality of Illumina and Nanopore raw reads were examined with fastqc (https://www.bioinformatics.babraham.ac.uk/projects/fastqc/) and nanopack [84]. PacBio Sequel II lacked a phred score. GC content in each sample was calculated using the gecee program from the EMBOSS package [85]. Illumina raw reads were trimmed with Trimmomatic [86] and assembled using IDBA-UD [87]. To improve the quality of the PacBio reads, we generated Highly Accurate Single-Molecule Consensus Reads (CCS reads) using the CCS program of the SMRT-link package. The minimum number of full-length subreads required to generate a CCS read was set to 5, 10 and 15 (99, 99.9 and 99.95 base call accuracy, respectively. Nanopore and PacBio (raw and CCS reads) were assembled using the following assemblers: SPAdes [88] with the metagenome option and performing a co-assembly with the Illumina trimmed reads; Flye [89] with the metagenome option; and HiCanu [90]. MetaFlye is a de novo assembler that follows the classical de Bruijn graphs, although it allows for approximate sequence matches. Canu, on the other hand, applies an overlapping strategy for de novo assembly. Lastly, SPAdes needs both short-reads and long-reads to perform a hybrid assembly. However, in the latter case, LR are only used for gap closure and repeat resolution. Given that the error-rate in PacBio reads can be significantly improved, we used the resulting CCS15 reads as single reads in the hybrid assembly with SPAdes, and in that case, CCS reads can be used together with the Illumina reads for graph construction, gap closure and repeat resolution.

### Taxonomic and Functional annotation of PacBio reads and assemblies

Prodigal [91] was used to predict genes from the assembled contigs retrieved from the individual assemblies of Illumina, Nanopore, and PacBio reads, as well as from the PacBio CCS reads. tRNA and rRNA genes were predicted using tRNAscan-SE [92] and barrnap (https://github.com/tseemann/barrnap), respectively. Predicted protein-encoded genes were taxonomical and functional annotated against the NCBI NR database using DIAMOND [93] and against COG [94] and TIGRFAM [95] using HMMscan [96].

### Taxonomic classification of metagenomic reads

16S rRNA gene sequences were retrieved from Illumina and PacBio reads. Candidate Illumina sequences in a subset of 20 million reads were extracted using USEARCH [97] after an alignment against a non-redundant version of the SILVA database [98]. Sequences that matched to this database with an E-value < 10^−5^ were considered potential 16S rRNA gene fragments. Then, ssu-align was used to identify true sequences aligning these candidate sequences against archaeal and bacterial 16S rRNA hidden Markov models (HMM). For the long-read sequences, candidate 16S rRNA sequences were extracted using barrnap from total PacBio CCS15 reads. The resulting 16S rRNA sequences were classified using the sina algorithm [99] according to the SILVA taxonomy database. Illumina sequences were only classified if the sequence identity was ≥ 80% and the alignment length ≥ 90 bp. Sequences failing these thresholds were discarded.

In addition, a total of 170 CCS15 contigs containing 16S and 23S rRNA genes of the phylum Cyanobacteria were selected to perform an internal transcribed spacer (ITS) phylogenetic tree, using the maximum-likelihood approach in iqtree [100], with 1000 bootstraps and the Jukes-Cantor model of substitution. Reference cyanobacterial ITS sequences were downloaded from the NCBI database.

### Genome reconstruction

Assembled contigs longer than 5Kb were assigned to a phyla classification if at least 50% of the genes shared the same best-hit taxonomy. Contigs failing this threshold were grouped as unclassified. To bin the contigs into MAGs, their taxonomic affiliation (including the unclassified) was used together with the principal component analysis of tetranucleotide frequencies, GC content, and coverage values within this sample and several metagenomic samples described in previous studies from the Mediterranean Sea [31,33,34,83]. Tetranucleotide frequencies were computed using wordfreq program in the EMBOSS package, and the principal component analysis was performed using the FactoMineR package [101]. Coverage values were calculated by the alignment of metagenomic reads (in subsets of 20 million reads) against contigs using BLASTN [102] (99% identity, > 50 bp alignment). Reads were normalized by the size of the contig in Kb and by the size of the metagenome in Gb (RPKGs). The degree of completeness and contamination of the resulting MAGs were estimated using CheckM [103]. Average nucleotide identity (ANI) between MAGs and the reference genome was calculated using JSpecies software with default parameters [104].

### Retrieval of relevant genes from the assemblies and the PacBio CCS reads

Predicted protein sequences of contigs and PacBio CCS15 reads longer than 5 Kb were compared against several downloaded and custom datasets. HMM were constructed using hmmbuild [96] for two curated databases of type-1 and type-3 rhodopsins. Searches were performed using hmmscan and only hits with a E-value < 10^−20^ were considered. To remove redundant proteins, sequences were clustered from 100 to 30% identity using cd-hit [105]. Glycosyl transferases (GT) were retrieved using dbCAN [106] against the Carbohydrate-Active enZYmes (CAZy) database [107]. To consider GTs involved in the flexible genome, only genomic fragments with ≥ 5 GTs and E-values < 10^−40^ were analyzed.

Lastly, the bacterial version of the secondary metabolite biosynthesis database (antiSMASH) was used to identify and classify [108] polyketide synthases (PKS) gene clusters from contigs and PacBio CCS15 reads longer than 5 Kb, and their taxonomic affiliation was based on consensus, that is > 70% of proteins encoded in a contig should share the same taxonomy (see above).

### Recovery and annotation of novel CRISPR-Cas systems

Sequences ≥ 5Kb long were screened using CRISPR-detect [69] and CRISPR-cas finder [109] tools. Only sequences matching in both methods and with an evidence value ≥ 3 were kept. The taxonomical affiliation of CCS reads and assembled contigs was based on the annotation of coded proteins (>70 % must share the same taxon). To find the putative target, CRISPR spacers were aligned using the blastn-short algorithm against nearly 200,000 phages collected and classified in [110]. Only matches with >97% identity and 100% alignment were considered. We also expanded the search including numerous metagenomic and viromic assemblies recovered from the Mediterranean Sea [31,33,34,83] and other marine samples [30,53].

### Data availability

Metagenomic datasets have been submitted to NCBI SRA and are available under BioProject accession number PRJNA674982 (Illumina reads: MedWinter-FEB2019-I [SAMN16686071]; Nanopore reads: MedWinter-FEB2019-NP [SAMN16686072]; and PacBio CCS reads: MedWinter-FEB2019-PBCCS15 [SAMN16686073]).

## ACKNOWLEDGEMENTS

This work was supported by grants “VIREVO” CGL2016-76273-P [AEI/FEDER, EU] (cofounded with FEDER funds) from the Spanish Ministerio de Economía, Industria y Competitividad, “HIDRAS3” PROMETEU/2019/009 from Generalitat Valenciana and 5top100-program of the Ministry for Science and Education of Russia to FRV. JHM was supported by a PhD fellowship from the Spanish Ministerio de Economia y Competitividad (BES-2014-067828). MLP was supported by a postdoctoral fellowship from the Spanish Ministerio de Economía, Industria y Competitividad (IJCI-2017-34002).

## AUTHORS’ CONTRIBUTIONS

The work was conceived by FRV. The analysis was carried out by JHM, MLP, and FRV. The manuscript was written by FRV, JHM, and MLP. All authors read and approved the final version.

## SUPPLEMENTARY MATERIAL

**Additional File 1: Figure S1**. Summary of the base-call accuracy, converted from the phred score, of the Nanopore raw reads. Upper and right histograms represent length and base-call accuracy distributions, respectively. **Figure S2**. Histogram representation of the GC content of the three sequencing methods: Illumina (black), Nanopore (red) and PacBio (blue). The resulting GC content after read correction of LR samples are also represented as dashed lines. **Table S1**. Relative abundance of 16S rRNA reads, based on the SILVA database. **Table S2**. Summary statistics of the assembly of PacBio raw reads using three assemblers (metaSPAdes, metaFlye and CanuHiFi) and applying different subsets of reads. **Table S3**. Genome parameters of MAGs recovered in this study with ANI > 99.5% to MAGs retrieved from the same sampling site in the Mediterranean Sea.

