## Additional File 1 for "Long read metagenomics, the next step?"

Evolutionary Genomics Group, División de Microbiología, Universidad Miguel Hernández, Apartado 18, San Juan de Alicante, 03550 Alicante, Spain.

### SUPPORTING INFORMATION

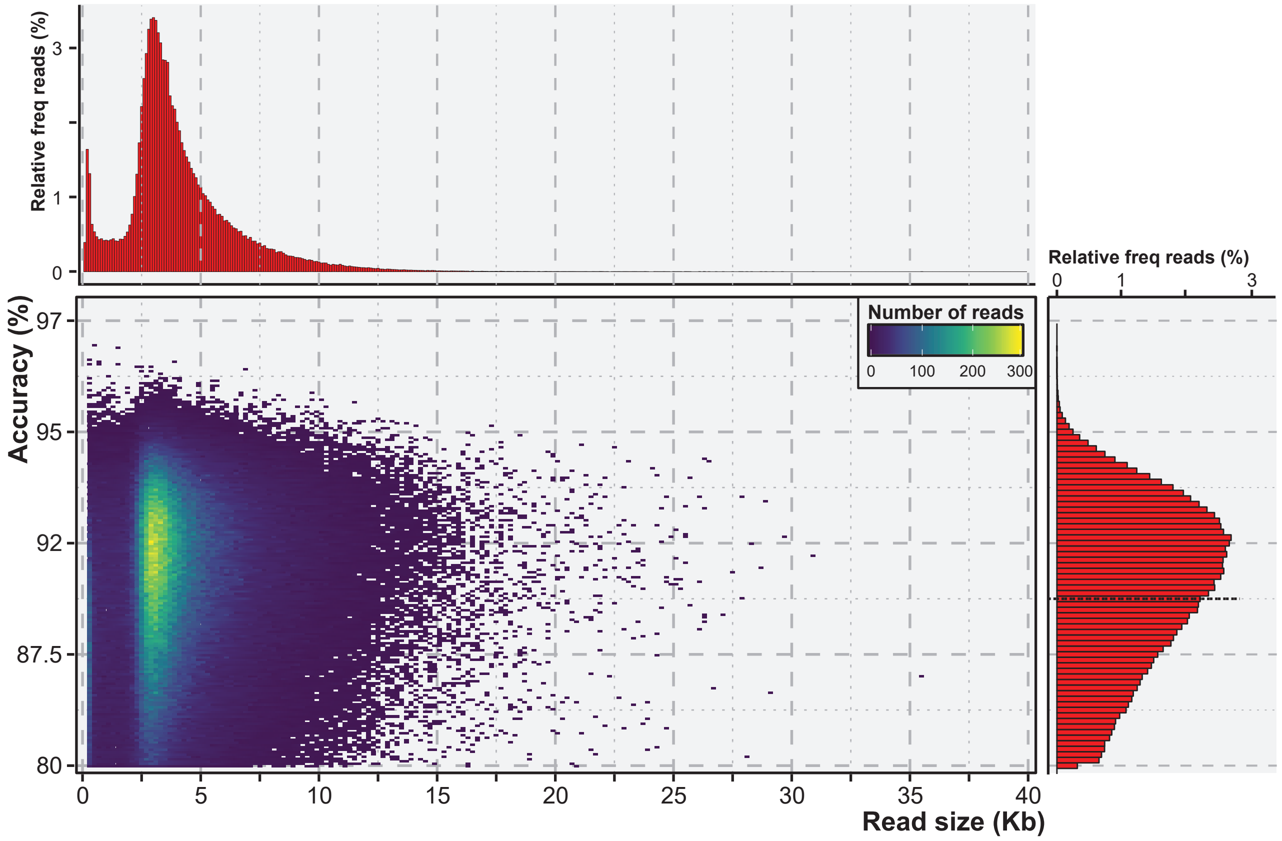


**Figure S1.** Summary of the base-call accuracy, converted from the phred score, of the Nanopore raw reads. Upper and right histograms represent length and base-call accuracy distributions, respectively.

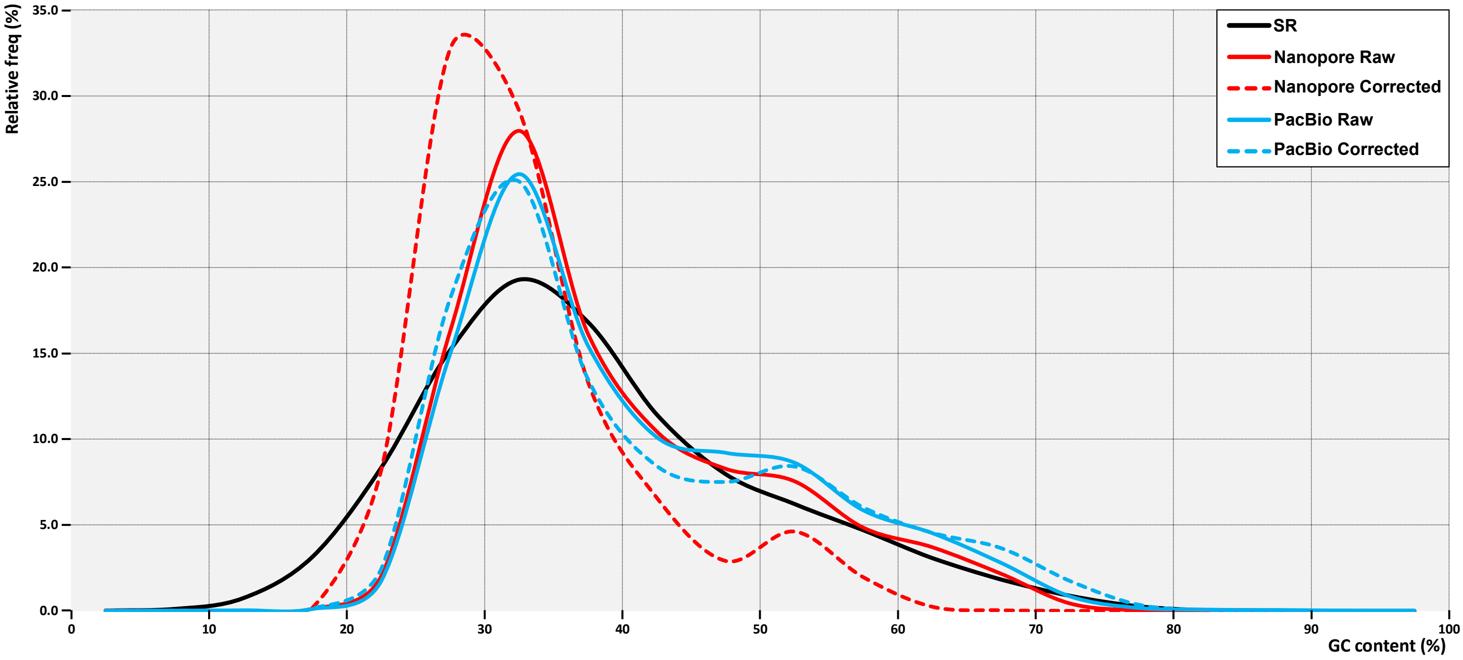


**Figure S2.** Histogram representation of the GC content of the three sequencing methods: Illumina (black), Nanopore (red) and PacBio (blue). The resulting GC content after read correction of LR samples are also represented as dashed lines.

**Table S1. Relative abundance of 16S rRNA reads.**

| Phylum | Group | SR (%) | LR CCS15 (%) |
|---|---|---|---|
| Euryarchaeota | | 2.8 | 2.9 |
| | Marine Group II | 2.6 | 2.7 |
| | Marine Group III | 0.1 | 0.1 |
| | Unclassified | 0.2 | 0.0 |
| Thaumarchaeota | | 1.1 | 1.6 |
| | Candidatus Nitrosopelagicus | 0.5 | 0.3 |
| | Candidatus Nitrosopumilus | 0.5 | 1.3 |
| | Unclassified | 0.1 | 0.0 |
| Actinobacteriota | | 2.3 | 2.3 |
| | Candidatus Actinomarina | 1.5 | 1.5 |
| | Microtrichales (Med-Acidi like) | 0.4 | 0.8 |
| | Unclassified | 0.3 | 0.0 |
| Bacteroidota | | 6.7 | 6.2 |
| | Cytophagales | 0.4 | 0.4 |
| | Flavobacteriales | 6.0 | 5.8 |
| | Unclassified | 0.3 | 0.0 |
| Chloroflexi | | 0.8 | 0.7 |
| | SAR202 clade | 0.7 | 0.6 |
| | JG30-KF-CM66 | 0.1 | 0.1 |
| Cyanobacteria | | 9.1 | 6.9 |
| | Prochlorococcus | 1.1 | 2.0 |
| | Synechococcus | 5.1 | 4.9 |
| | Other Groups | 2.9 | 0.0 |
| Dadabacteria | | 0.2 | 0.4 |
| Marinimicrobia (SAR406 clade) | | 1.8 | 2.3 |
| Nitrospinota | | 0.2 | 0.2 |
| Planctomycetota | | 0.7 | 0.4 |
| | Phycisphaerales | 0.1 | 0.1 |
| | Pirellulales | 0.4 | 0.3 |
| | Unclassified | 0.2 | 0.0 |
| Alphaproteobacteria | | 47.7 | 49.9 |
| | Defluviicoccales | 0.5 | 0.4 |
| | OCS116 clade | 0.9 | 1.3 |
| | PS1 clade | 0.1 | 0.1 |
| | SAR116 clade | 2.3 | 2.5 |
| | Rhizobiales | 0.4 | 0.0 |
| | Rhodobacterales | 3.0 | 2.6 |
| | Rhodospirillales | 4.0 | 4.3 |
| | SAR11 clade | 34.7 | 37.4 |
| | Thalassobaculales | 0.2 | 0.4 |
| | Unclassified | 1.6 | 0.5 |
| Gammaproteobacteria | | 20.4 | 20.5 |
| | Alteromonadales | 0.4 | 0.1 |
| | Burkholderiales | 0.4 | 0.3 |
| | OM60(NOR5) clade | 0.9 | 1.0 |
| | SAR92 clade | 1.1 | 0.9 |
| | Ectothiorhodospirales | 0.8 | 1.2 |
| | HOC36 | 0.3 | 0.2 |
| | KI89A clade | 0.4 | 0.6 |
| | Oceanospirillales | 1.1 | 0.4 |
| | OM182 clade | 0.2 | 0.3 |
| | SAR86 clade | 10.2 | 12.0 |
| | Steroidobacterales | 0.4 | 0.4 |
| | SUP05 cluster | 1.0 | 1.2 |
| | Thiotrichales | 0.5 | 0.3 |
| | UBA10353 marine group | 0.2 | 0.2 |
| | Unclassified | 1.8 | 0.8 |
| SAR324 clade(Marine group B) | | 0.8 | 1.2 |
| Verrucomicrobiota | | 3.5 | 3.5 |
| | Kiritimatiellales | 0.1 | 0.1 |
| | Arctic97B-4 marine group | 0.4 | 0.4 |
| | Opitutales | 2.2 | 2.3 |
| | Pedosphaerales | 0.2 | 0.2 |
| | Verrucomicrobiales | 0.4 | 0.4 |
| Other Groups (< 0.2%) | | 0.6 | 0.8 |
| Unclassified | | 1.3 | 0.4 |

**Table S2. Summary statistics of the assembly of PacBio raw reads with three different assemblers at five sequencing depths.**

| | metaSPAdes | | | | | metaFlye | | | | | Canu | | | | |
|---|---|---|---|---|---|---|---|---|---|---|---|---|---|---|---|
| **PacBio reads (Gb)** | **100K (1.1)** | **500K (5.6)** | **1M (11.3)** | **5M (56.6)** | **10M (113.7)** | **100K (1.1)** | **500K (5.6)** | **1M (11.3)** | **5M (56.6)** | **10M (113.7)** | **100K (1.1)** | **500K (5.6)** | **1M (11.3)** | **5M (56.6)** | **10M (113.7)** |
| **Number of Contigs** | 12,886 | 16,872 | 20,138 | 30,303 | 33,362 | 25 | 497 | 1,488 | 11,044 | 23,152 | 159 | 6,417 | 13,304 | 49,681 | 122,210 |
| **Assembly Size (Mb)** | 127.13 | 168.40 | 203.42 | 328.86 | 379.24 | 0.62 | 14.79 | 44.69 | 341.75 | 707.47 | 1.3 | 59.6 | 134.7 | 582.4 | 1435.7 |
| **Largest Contig Size (Mb)** | 0.20 | 0.20 | 0.21 | 0.35 | 0.37 | 0.04 | 0.13 | 0.29 | 0.99 | 2.22 | 0.04 | 0.24 | 0.27 | 0.85 | 0.57 |
| **Average Contig Size (Kb)** | 9.9 | 10.0 | 10.1 | 10.9 | 11.4 | 24.8 | 29.8 | 30.0 | 30.9 | 30.6 | 8.5 | 9.3 | 10.1 | 11.7 | 11.7 |
| **Number of Proteins** | 137,824 | 190,325 | 234,948 | 425,304 | 473,163 | 429 | 9,776 | 29,768 | 218,880 | 460,967 | 2,399 | 84,899 | 179,347 | 748,258 | 1,905,216 |
| **Average Protein Size (aa)** | 264.8 | 245.5 | 235.5 | 213.9 | 204.9 | 79.4 | 77.5 | 76.2 | 76.4 | 78.2 | 114.1 | 179.2 | 196.9 | 207.2 | 199.5 |
| **Proteins / Mb** | 1084.1 | 1130.2 | 1155.0 | 1293.3 | 1247.7 | 691.9 | 661.1 | 666.1 | 640.5 | 651.6 | 1782.8 | 1424.3 | 1331.8 | 1284.8 | 1327.0 |

**Table S3. Genome parameters of MAGs recovered in this study with ANI > 99.5% to MAGs retrieved from the same sampling site in the Mediterranean Sea.**

| Genome | Metagenome | Genome Size (bp) | #Contigs | Largest Contig Size (bp) | Average Contig Size (bp) | Completeness (%) | Contamination (%) |
|---|---|---|---|---|---|---|---|
| PS1 MED-G09 (GCA_002457395.1) | a | 746,185 | 18 | 121,164 | 41,454.7 | 56.6 | 0.0 |
| | LR CCS15 | 1,497,191 | 3 | 1,091,529 | 499,063.7 | 89.5 | 1.2 |
| Rhodobacteraceae MED-G07 (GCA_002457115.1) | a | 1,097,455 | 41 | 95,239 | 26,767.2 | 56.6 | 0.0 |
| | LR CCS15 | 2,430,758 | 21 | 387,711 | 115,750.4 | 85.5 | 0.8 |
| Alphaproteobacteria MED-G51 (GCA_003331375.1) | a | 1,161,306 | 38 | 124,460 | 30,560.7 | 68.8 | 0.0 |
| | LR CCS15 | 1,691,115 | 10 | 364,231 | 169,111.5 | 86.0 | 0.0 |
| Rhodobacteraceae MED-G52 (GCA_003332035.1) | a | 1,390,107 | 48 | 174,535 | 28,960.6 | 62.7 | 0.0 |
| | LR CCS15 | 2,439,284 | 12 | 552,512 | 203,273.7 | 90.8 | 0.7 |
| Rhodobacteraceae MED-G111 (GCA_004213335.1) | b | 1,869,508 | 56 | 171,379 | 33,384.1 | 72.4 | 0.0 |
| | LR CCS15 | 2,278,673 | 20 | 359,451 | 113,933.6 | 86.9 | 0.0 |
| Rhodobacteraceae MED-G112 (GCA_004213125.1) | b | 1,698,068 | 187 | 31,943 | 9,080.6 | 56.0 | 0.9 |
| | LR CCS15 | 1,766,722 | 58 | 79,409 | 30,460.7 | 56.7 | 0.1 |
| Cryomorphaceae MED-G11 (GCA_002457075.1) | a | 851,326 | 30 | 130,165 | 28,377.5 | 73.9 | 0.0 |
| | LR CCS15 | 1,221,852 | 11 | 210,260 | 111,077.5 | 86.4 | 2.8 |
| Rhodothermaeota MED-G16 (GCA_002457035.1) | a | 1,015,913 | 39 | 95,123 | 26,049.1 | 54.9 | 0.6 |
| | LR CCS15 | 1,643,288 | 19 | 206,509 | 86,488.8 | 82.4 | 1.1 |
| Cryomorphaceae MED-G61 (GCA_003331885.1) | a | 870,081 | 36 | 97,093 | 24,168.9 | 66.9 | 0.0 |
| | LR CCS15 | 1,538,260 | 13 | 381,277 | 118,327.7 | 96.3 | 0.7 |
| Chloroflexi MED-G130 (GCA_004214105.1) | b | 1,310,007 | 9 | 487,332 | 145,556.3 | 88.1 | 0.0 |
| | LR CCS15 | 1,141,320 | 14 | 189,172 | 81,522.9 | 81.2 | 0.0 |
| Chloroflexi MED-G131 (GCA_004213445.1) | b | 1,010,725 | 43 | 91,403 | 23,505.2 | 78.7 | 0.0 |
| | LR CCS15 | 1,460,397 | 16 | 219,892 | 91,274.8 | 93.1 | 0.5 |
| Prochlorococcus MED-G72 (GCA_003331725.1) | a | 1,278,176 | 22 | 243,092 | 58,098.9 | 79.4 | 0.0 |
| | LR CCS15 | 1,615,769 | 8 | 607,950 | 201,971.1 | 98.4 | 0.0 |
| Prochlorococcus MED-G73 (GCA_003331715.1) | a | 781,345 | 37 | 59,189 | 21,117.4 | 55.8 | 0.0 |
| | LR CCS15 | 1,623,167 | 13 | 491,601 | 124,859.0 | 96.5 | 0.6 |
| Synechococcus MED-G67 (GCA_003331795.1) | a | 1,679,931 | 30 | 351,581 | 55,997.7 | 81.5 | 0.3 |
| | LR CCS15 | 1,556,683 | 41 | 108,381 | 37,967.9 | 71.8 | 0.8 |
| EUII MED-G37 (GCA_002457555.1) | a | 1,284,027 | 25 | 198,151 | 51,361.1 | 71.7 | 0.0 |
| | LR CCS15 | 1,008,693 | 29 | 86,510 | 34,782.5 | 49.0 | 0.0 |
| EUII MED-G38 (GCA_002457145.1) | a | 1,366,214 | 21 | 164,554 | 65,057.8 | 73.9 | 0.0 |
| | LR CCS15 | 1,155,561 | 26 | 158,711 | 44,444.7 | 65.9 | 0.0 |
| SUP05 MED-G23 (GCA_002456985.1) | a | 847,715 | 36 | 78,958 | 23,611.1 | 62.6 | 0.0 |
| | LR CCS15 | 913,584 | 33 | 51,158 | 27,684.4 | 19.0 | 0.0 |
| OM60/NOR5 MED-G26 (GCA_002456955.1) | a | 1,280,660 | 62 | 70,617 | 20,655.8 | 55.2 | 0.0 |
| | LR CCS15 | 2,599,316 | 5 | 1,501,514 | 519,863.2 | 95.2 | 0.0 |
| OM182 MED-G28 (GCA_002457215.1) | a | 2,947,553 | 25 | 413,588 | 117,902.1 | 88.7 | 0.5 |
| | LR CCS15 | 1,329,907 | 44 | 69,957 | 30,225.2 | 33.5 | 0.0 |
| SAR92 MED-G29 (GCA_002457245.1) | a | 1,295,755 | 53 | 96,029 | 24,448.2 | 71.1 | 0.0 |
| | LR CCS15 | 2,260,979 | 15 | 361,335 | 150,731.9 | 85.4 | 3.0 |
| Gammaproteobacteria MED-G80 (GCA_003331585.1) | a | 1,314,669 | 29 | 141,446 | 45,333.4 | 63.0 | 1.4 |
| | LR CCS15 | 1,757,721 | 7 | 540,954 | 251,103.0 | 79.6 | 2.1 |
| Gammaproteobacteria MED-G143 (GCA_004213285.1) | b | 676,909 | 56 | 31,337 | 12,087.7 | 46.8 | 0.4 |
| | LR CCS15 | 687,601 | 6 | 168,312 | 114,600.2 | 62.1 | 0.0 |
| Gammaproteobacteria MED-G148 (GCA_004213855.1) | b | 732,908 | 70 | 36,265 | 39.7 | 44.6 | 0.6 |
| | LR CCS15 | 1,639,117 | 5 | 587,642 | 327,823.4 | 78.5 | 1.7 |
| Bacteria MED-G45 (GCA_003332085.1) | a | 452,526 | 22 | 45,843 | 20,569.4 | 45.9 | 0.0 |
| | LR CCS15 | 984,988 | 9 | 336,464 | 109,443.1 | 89.0 | 2.2 |
| Bacteria MED-G46 (GCA_003331415.1) | a | 1,231,823 | 24 | 192,181 | 51,250.0 | 68.1 | 0.0 |
| | LR CCS15 | 1,661,099 | 11 | 531,357 | 151,009.0 | 93.4 | 1.1 |
| Bacteria MED-G176 (GCA_004321715.1) | b | 1,630,357 | 46 | 131,927 | 35,442.5 | 62.1 | 0.0 |
| | LR CCS15 | 905,771 | 26 | 77,504 | 34,837.3 | 32.8 | 0.0 |
| Phycisphaeraceae MED-G179 (GCA_004213685.1) | b | 2,107,155 | 124 | 85,114 | 16,993.2 | 72.0 | 0.0 |
| | LR CCS15 | 3,154,705 | 15 | 588,530 | 210,313.7 | 94.2 | 1.2 |
| Pedosphaeraceae MED-G185 (GCA_004321855.1) | b | 3,048,501 | 188 | 103,876 | 16,215.4 | 75.4 | 0.7 |
| | LR CCS15 | 4,156,947 | 66 | 215,054 | 62,984.0 | 87.8 | 1.7 |
| Pedosphaeraceae MED-G186 (GCA_004213515.1) | b | 2,735,641 | 103 | 169,050 | 26,559.6 | 77.9 | 0.0 |
| | LR CCS15 | 3,183,644 | 21 | 499,935 | 151,602.1 | 93.6 | 1.4 |
| Pedosphaeraceae MED-G187 (GCA_004213675.1) | b | 1,080,818 | 117 | 38,866 | 9,237.8 | 43.0 | 0.0 |
| | LR CCS15 | 3,083,832 | 48 | 345,717 | 64,246.5 | 78.7 | 1.3 |
| Puniceicoccaceae MED-G32 (GCA_002457235.1) | a | 671,118 | 30 | 47,853 | 22,333.3 | 69.6 | 0.0 |
| | LR CCS15 | 1,011,612 | 21 | 170,873 | 48,172.0 | 76.5 | 0.7 |
| **Averages** | **a-b** | **1,337,564** | **54** | **139,334** | **35,877** | **66** | **0.2** |
| | **LR CCS15** | **1,932,065** | **19** | **434,047** | **162,207** | **83** | **1.0** |

MAGs from [a]Haro-Moreno et al. 2018 and [b]Haro-Moreno et al. 2019
